# Characterizing concentration-dependent neural dynamics of 4-aminopyridine-induced epileptiform activity

**DOI:** 10.1101/252726

**Authors:** Timothy L Myers, Oscar C González, Jacob B Stein, Maxim Bazhenov

## Abstract

Epilepsy remains one of the most common neurological disorders. In patients, it is characterized by unprovoked, spontaneous, and recurring seizures or ictal events. Typically, inter-ictal events or large bouts of population level activity can be measured between seizures and are generally asymptomatic. Decades of research has focused on understanding the mechanisms leading to the development of seizure-like activity using various proconvulsive pharmacological agents, including 4-aimnopyridine (4AP). However, the lack of consistency in the concentrations used for studying 4AP-induced epileptiform activity in animal models may give rise to differences in the results and interpretation thereof. Indeed, the range of 4AP concentration in both *in vivo* and *in vitro* studies varies from 3μM to 40mM. Here, we explored the effects of various 4AP concentrations on the development and characteristics of hippocampal epileptiform activity in acute mouse brain slices of either sex. Using multielectrode array recordings, we show that 4AP induces hippocampal epileptiform activity for broad range of concentrations. The frequency component and the spatio-temporal patterns of the epileptiform activity revealed a dose-dependent response. Finally, in the presence of 4AP, reduction of KCC2 co-transporter activity by KCC2 antagonist VU0240551 prevented the manifestation of the frequency component differences between different concentrations of 4AP. Overall, the study predicts that different concentrations of 4AP can result in the different mechanisms behind hippocampal epileptiform activity, of which some are dependent on the KCC2 co-transporter function.

## Introduction

Potassium (K^+^) channel activation has been shown to be responsible for the repolarization of the membrane potential following the depolarization phase of an action potential (1-5). Differences in the type of K^+^-channels expressed in a given neuron influence the properties of action potentials produced by the neuron (1, 4). Outward going K^+^ A-type currents (I_A_) have been shown to regulate the action potential firing rate by modulating the inter-spike interval in response to subthreshold current injections (2-5). The I_A_ is mediated by multimeric channels comprised of mainly K_v_4 family α-subunits in combination with modulatory β-subunits (6). Functional knock-out, through expression of dominant negative K_v_4.2 α-subunits (K_v_4.2DN) in rat cortical pyramidal neurons has been shown to selectively eliminate I_A_, resulting in reduced action potential thresholds, increased action potential duration, and increased neuronal excitability Functional knock-out, through expression of dominant negative K_v_4.2 (5). Additionally, experiments in rat hippocampal neurons have shown that nearly all of the K^+^ current underlying the repolarization phase of an action potential can be accounted for by the I_A_ and the dendrotoxin-sensitive K^+^ D-current (I_D_) (3). Due to its important role in regulating neuronal excitability, it is not surprising that the I_A_ has been the subject of intense study in its relation to epileptic seizures.

Certain genetic epilepsies have been linked to misregulated or mutated K^+^ channels in humans (7-14). Some patients suffering from intractable temporal lobe epilepsy have been shown to express K_v_4 α-subunit mutations, specifically truncation mutations in K_v_4.2 α-subunits (8, 14). In these patients, the mutation manifests as a reduction of the I_A_ thereby leading to hyperexcitability. Similarly, I_A_ and I_D_ antagonist 4-aminopyridine (4AP) has been shown, in *in vivo* and *in vitro* studies, to cause hyperexcitability and the development of seizure-like epileptiform discharges (15-23). Recent work suggests that the mode of action of 4AP in epileptiform onset is not through its direct reduction of I_A_ or I_D_ and increased principle neuron excitation; rather, 4AP increases the excitability of both inhibitory and excitatory neurons, the former of which is pivotal in the development of seizure-like activity (8, 14). In these patients, the mutation manifests as a reduction of the I_A_ thereby leading to hyperexcitability. Similarly, I_A_ and I_D_ antagonist 4-aminopyridine (4AP) has been shown, in *in vivo* and *in vitro* studies, to cause hyperexcitability and the development of seizure-like epileptiform discharges (15, 17-19, 21, 23, 24). Previous studies have shown that the potassium-chloride co-transporter isoform 2 (KCC2) plays an important role in the generation of 4AP-induced seizure (15, 17-19, 22). Indeed, in rat brain slices treated with 4AP, reduction of KCC2 activity prevents seizure-like discharges while interictal activity remained relatively intact (18). The mechanism and extent to which KCC2 influences the properties of 4AP-induced inter-ictal activity remains to be fully understood.

Previous studies have attempted to fill this gap in our understanding precise effects of 4AP on neuronal activity, but lack of consistency in the methods and concentrations of 4AP, ranging from 3 μM to 40 mM, used for such studies prevents the development of a complete picture (18, 20, 21, 23, 25-28). In our new study, we explored the concentration-dependent effects of 4AP on the spatiotemporal properties of induced epileptiform activity in acute mouse hippocampal brain slices. We found that bath applied 4AP produced dose-specific epileptiform bursts, which properties - frequency and spatial pattern - depended on the 4AP concentration. Additionally, the reduction of KCC2 co-transporter activity through bath applied VU0240551 prevented the generation of the higher frequency component of epileptiform bursts at high 4AP concentrations.

## Materials and Methods

### Animals

All experiments and procedures were performed according to University of California, Riverside Institutional Animal Care and Use Committee-approved protocols. Wild-type (Jackson Laboratory, C57BL/6J, stock number 000664) mouse colonies were bred and maintained in house in order to generate pups for this study. All mice were provided fresh water and mouse chow *ad libitum*, and a consistent circadian cycle was maintained by the vivarium facility. Both male and female mice were used in this study.

### Brain slice preparation

Postnatal day (P) 15-20 mice were anesthetized with isoflurane and quickly decapitated. The brain was rapidly removed and submerged in ice-cold, low [Ca^2+^]_o_, high sucrose dissecting solution continuously oxygenated with carbogen (95%-5% O_2_-CO_2_) gas. Horizontal whole brain slices (300 μm) were made with a Leica 1200S vibratome. Slices containing both hippocampus and entorhinal cortex were recovered in oxygenated normal artificial cerebrospinal fluid (ACSF) for 1 h at 32C, and stored at room temperature. The standard dissecting solution contained (mM) 87 NaCl, 2.5 KCl, 25 NaHCO_3_, 1.25 NaH_2_PO_4_, 4 MgCl_2_, 0.5 CaCl_2_, 10 D-glucose, 75 sucrose.

During recordings, brain slices were continuously perfused with ACSF (3 ml/min flow rate) containing (mM) 125 NaCl, 2.5 KCl, 25 NaHCO_3_, 1.25 NaH_2_PO_4_, 1 MgCl_2_, 2 CaCl_2_, 25 D-glucose, 10 sucrose. Both solutions were maintained at pH 7.4 by continuous oxygenation with carbogen gas mixture. All chemicals were obtained though Fisher Scientific unless otherwise specified.

### Electrophysiological recordings

Multielectrode array (MEA) recordings were performed on a 60-channel perforated array (60pMEA200/30-Ti) with a low-noise amplifier (MEA1060-BC) from MultiChannel Systems. Hippocampal slices were prepared, as described above, placed on the array, and positioned such that the CA1, CA3, and dentate gyrus (DG) were centered over the recording electrodes. Channels with high noise level were silenced prior to recording. Experiments consisted of an initial 30 min. ACSF control followed by eight consecutive 30 min. conditions of increasing 4-aminopyridine concentrations (25 – 200 μM). To test the role of the KCC2 co-transporter in the characteristics of 4AP-induced epileptiform activity, we followed the same procedure outlined above, but added 10 μM of the KCC2 co-transporter antagonist VU0240551 to the perfusate containing 4AP. All MEA recordings were performed at 32C, which was sufficient to generate stable epileptiform activity. For 4AP only experiments n = 4 slices from 4 different mice, and for 4AP + VU0240551 n = 6 slices from 6 different mice.

### Analysis

Data were acquired using MC Rack software (MultiChannel Systems) and exported to MATLAB (MathWorks) for further processing. Data from all 60 MEA channels were collected at 25 kHz and downsampled offline to 5 kHz. Synchronized network activity or epileptiform bursts recorded by the MEA were detected using our previously described percentile-based method (29). Briefly, a low (1.25 ± 0.75) and high (98.75 ± 0.75) percentile were used on unfiltered data from a single channel recording from each of the hippocampal subfields mentioned above. The low percentile was used for events with negative polarity, while the high percentile was used for events with positive polarity. The amplitude corresponding to the low or high percentile were used as a threshold for detecting the epileptiform bursts. Similar to (29), burst onset was defined as the moment when the LFP crossed the percentile-defined threshold.

The bursts where then extracted from all channels with 500 ms before and after burst onset, and a non-overlapping period of 1 s was used. Inter-event intervals were computed as the difference between timing of the peak of a given event and that of the event prior to it. Event frequency was computed as the inverse of the inter-event interval. Fourier transforms were performed on the unfiltered, downsampled data using Matlab function fft. We used a sliding window of size 60 s with a 30 s overlap to compute the Fourier transform. The mean was computed across all resulting transform signals across all slices for each condition. Data are presented as mean ± standard error (SEM).

## Results

We induced epileptiform activity throughout the hippocampal cortex by bath application of 4-aminopyridine (4AP). Local field potentials (LFPs) were recorded during epileptiform activity using a 60-channel MEA, from the hippocampal subfields CA1, CA3, and DG positioned over the MEA electrodes (fig 1A). Control, ACSF only recordings, showed a minimal multiunit activity (data not shown). Spontaneous epileptiform activity was not observed in control conditions. Bath applied 4AP at all concentrations tested in this study (25 – 200 μM) resulted in a transition from quiescence to a seizure-like state characterized by stereotyped continuous epileptiform activity (fig 1). Epileptiform activity was robust, and highly synchronized across all MEA electrodes (fig 1B). Single channel recordings from CA1, CA3, and DG revealed the bias towards CA3 as the generator of the largest epileptiform activity (fig 1C, E). Figure 1D shows all the extracted epileptiform bursts (see Methods) from a single channel in CA3 superimposed (black traces) with the average burst superimposed in red. This revealed that the events recorded from the same channel show similar temporal properties throughout the duration of the burst. The temporal similarities in burst progression were not constrained to recordings from CA3. Figure 1F shows heatmaps of the all bursts recorded in the same channels from CA1, CA3, and DG revealing stereotypes activity induced throughout the hippocampus by 4AP.

**Fig 1.**
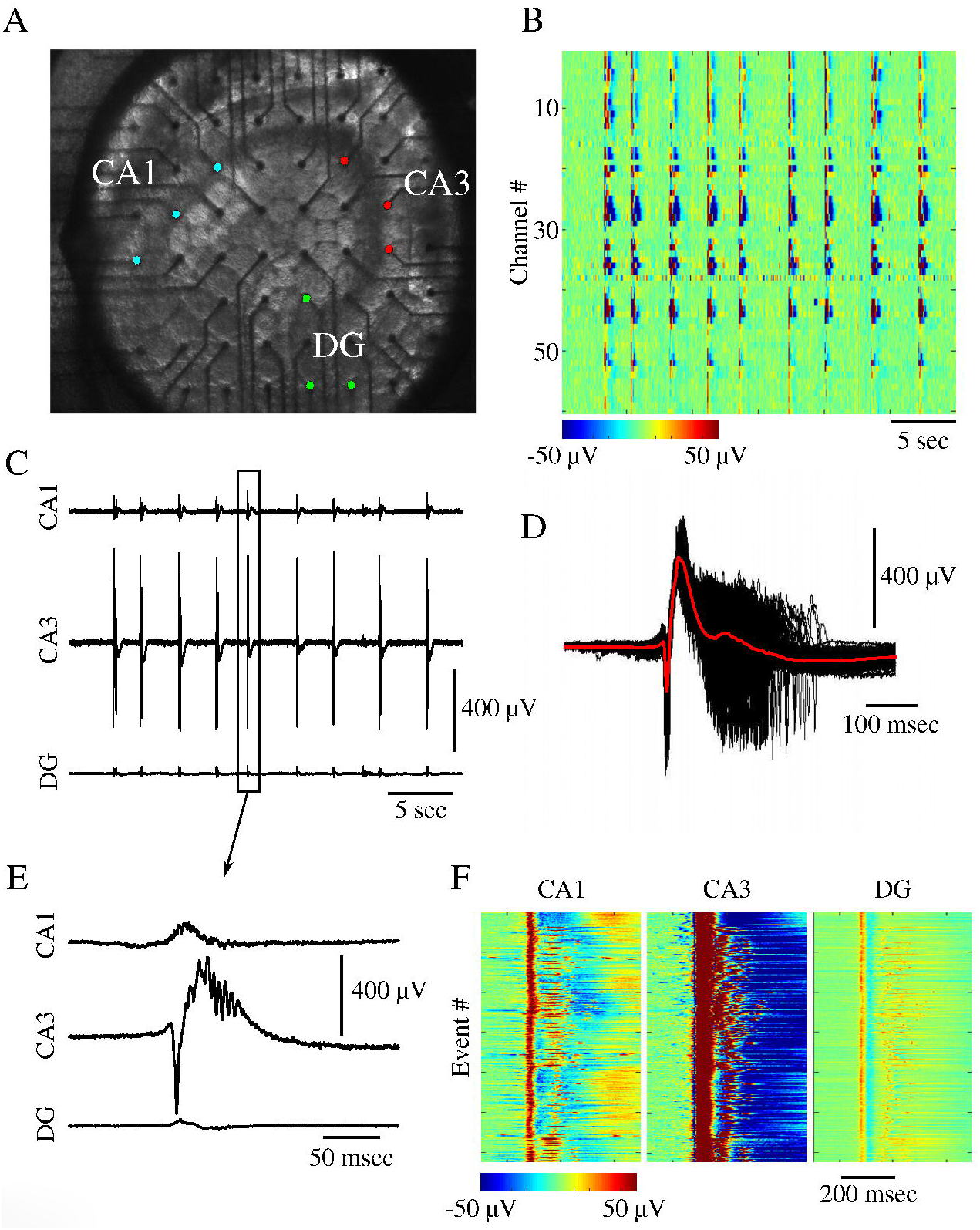
4AP elicits epileptiform activity across the mouse hippocampus. **A**, image showing the position of an acute mouse hippocampal brain slice on the multielectrode array (MEA). CA1, CA3, and dentate gyrus (DG) subfields have been labeled, and colored electrodes correspond to electrodes within the structures. **B,** epileptiform activity induced with 100 μM 4AP recorded by 60 channel MEA. Color scale represents voltage as indicated by scale bar. **C,** epileptiform bursts recorded from single channels within the CA1 (top), CA3 (middle), and DG (bottom). **D,** time aligned epileptiform bursts recorded from single CA3 electrode. The average signal is superimposed in red. **E,** zoom in on a single burst from C. **F,** all epileptiform bursts recorded from the same channels in C. As mentioned above, 100 μM 4AP resulted in stereotyped epileptiform activity (fig 1). Since 4AP selectively blocks outward going K^+^ A- and D-currents, we hypothesized that varying the concentration of 4AP may lead to differences in the properties of the resulting epileptiform activity. To test this, we varied the concentration of bath applied 4AP, ranging from 25 – 200 μM in 25 μM step sizes. Epileptiform activity was reliably elicited for all 4AP concentrations from that range. We first examined the differences in the spatio-temporal distribution of epileptiform bursts throughout the hippocampus. Figure 2 shows the propagation of a single epileptiform burst (such as, e.g., shown in fig 1D), resulting from different concentrations of bath applied 4AP, through the hippocampus. For all concentrations, the initiation of the epileptiform burst occurred near the CA3 subfield before propagating to the rest of the hippocampus (fig 2). These epileptiform bursts initiated with a positive electrical potential (source) within the areas of the proximal and distal dendrites, which was accompanied by a negative potential (sink) within stratum pyramidale. Similar findings have been reported for epileptiform activity induced through elevated [K^+^]_o_ (29). This was then followed by a reversal of the polarity of the field potentials in their respective locations.

**Fig 2.**
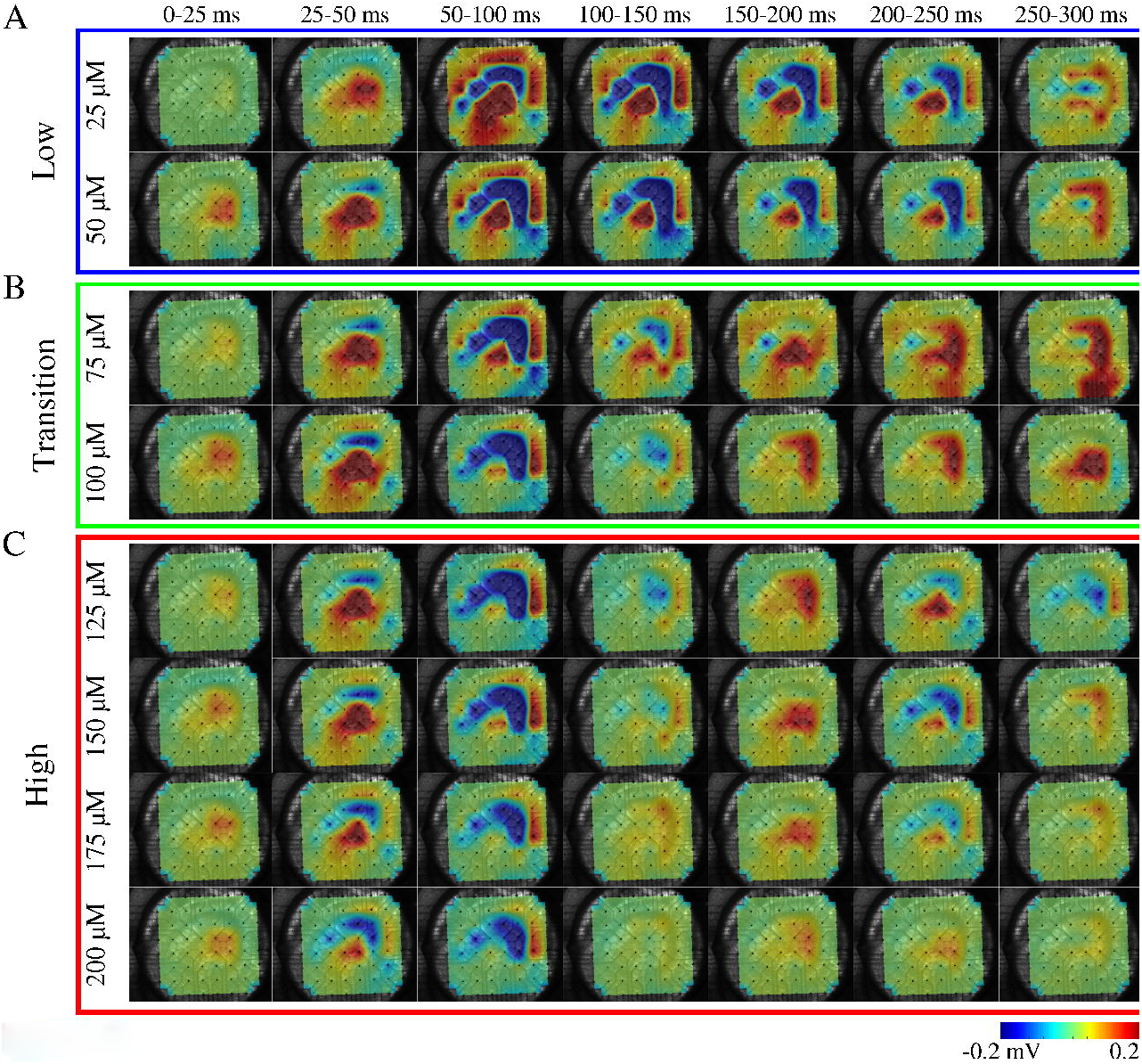
Concentration-dependent differences in spatiotemporal propagation of epileptiform activity. Sequence of averaged (25 ms) LFP activity for one event. Each row corresponds to a different 4AP concentration. **A**, low 4AP concentrations. **B,** transition concentrations. **C,** high concentrations. Color represents voltage as indicated by the colorbar. We observed a much wider spatial distribution of epileptiform activity for low concentrations (fig 2). As seen for 25 and 50 μM concentrations, the electrical activity recorded during the 50 – 100 ms frame showed electrical activity propagating from CA3 to CA1. As mentioned previously, the stratum pyramidale exhibited negative LFP while the regions containing pyramidal cell dendrites exhibited positive LFP during early and middle phase of the epileptic burst, which reversed in time during later phase of the event. Additionally, in low concentrations (25 – 50 μM) the negative potential recorded from stratum pyramidale remained for 200 ms before reversing polarity. Unlike the prolonged period of negative LFP for low concentrations of 4AP, concentrations ranging from 75 – 125 μM exhibited a slightly prolonged period of positive LFP within CA3 and DG (compare time frames 150-200ms, 200-250ms, and 250-300ms for concentrations 25 – 125 μM of figs 2A and 2B). For high concentrations of 4AP (150 – 200 μM), the spatial distribution of the epileptiform burst became more focal and short lived (fig 2, red box). Indeed, little activity was observed following 100 ms after the initiation of the epileptiform burst. These data suggest that the extent of the reduction of A- and D-current activity may cause substantial differences in the dynamics of the epileptiform activity. The differences in the spatio-temporal distributions of the epileptiform activity shown in figure 2 could reflect underlying differences in the frequency components of the epileptic bursts. To explore this, we computed Fourier transforms of the electrical signals generated by the hippocampal slice under the conditions of bath applied 4AP. We observed that low concentrations of 4AP, 25 – 50 μM, generated epileptiform bursts that were mainly characterized by low frequencies <5 Hz (fig 3A). The activity generated with 75 μM 4AP showed a small increase in higher frequencies (around 7 – 15 Hz), a trend which was also observed in epileptiform bursts generated at 100 μM 4AP (fig 3B). These higher frequencies became more prominent at higher concentrations, 125 – 200 μM (fig 3C). Insets show example of a single corresponding epileptiform burst for the concentration indicated showing increased burst complexity with increasing 4AP concentration. Interestingly, the low frequency component of the epileptiform bursts remained largely intact at high concentrations. These data, along with the previously described data (fig 2), support our hypothesis that differentially impairing A- and D-current activity results in different epileptiform activity profiles.

**Fig 3.**
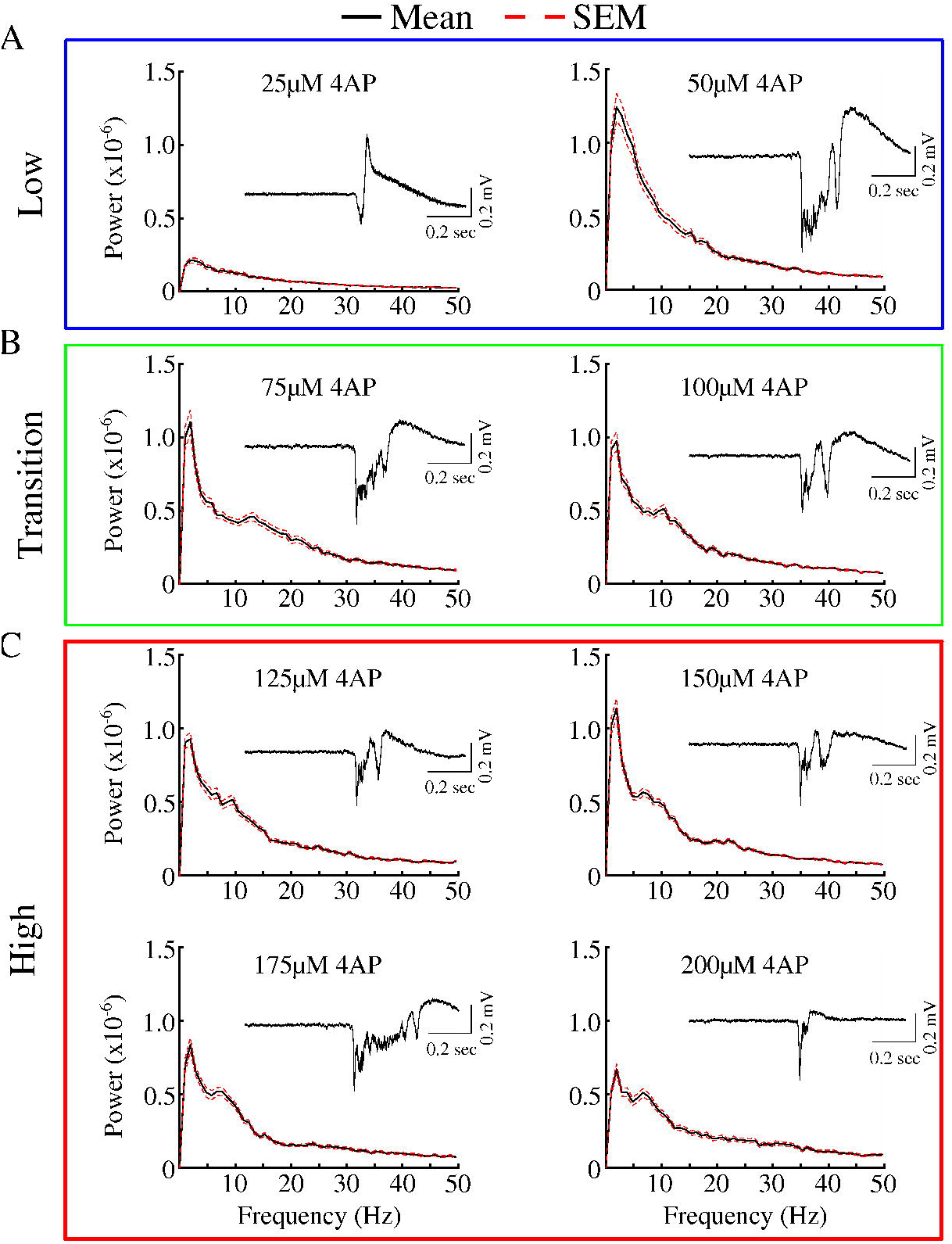
Concentration-dependent differences in epileptiform spectrograms. Averaged spectrograms of activity induced by different concentrations of 4AP. The mean is plotted in black solid line and standard error (SEM) in red broken line. **A,** low 4AP concentrations. **B,** transition concentrations. **C,** high concentrations. Insets are of single representative epileptiform bursts elicited by the corresponding 4AP concentration. Recent evidence has suggested that the potassium-chloride co-transporter isoform 2 (KCC2) may play a key role in the generation of 4AP-induced epileptiform activity (15, 17-19, 22). We hypothesized that the reduction of KCC2 activity by KCC2 antagonist VU0240551 may reduce the complexity of epileptiform activity elicited at higher concentrations of 4AP, resulting in more stereotyped epileptiform activity across various 4AP concentrations. To test this hypothesis, we bath applied KCC2 specific antagonist VU0240551, and varied the bath applied 4AP concentrations. Similar to our previous experiment (fig 1), we observed 4AP-induced epileptiform for all concentrations tested with CA3 exhibiting the largest amplitude epileptiform bursts. Low 4AP concentration conditions (25 – 50 μM), once again, produced epileptiform activity dominated by low frequency activity (fig 4A). Interestingly, increasing the 4AP concentrations from 75 – 200 μM did not led to the power spectrum peak at higher frequencies as seen previously in conditions of intact KCC2 (compare figs 3B,C and 4B,C). Additionally, the epileptiform activity elicited in the presence of the KCC2 antagonist became stereotyped across 4AP concentrations (compare fig 4 insets). These finding suggest that the high frequency component of the epileptiform activity for high 4AP concentrations may be dependent on the recruitment of KCC2-dependent mechanisms.

**Fig 4.**
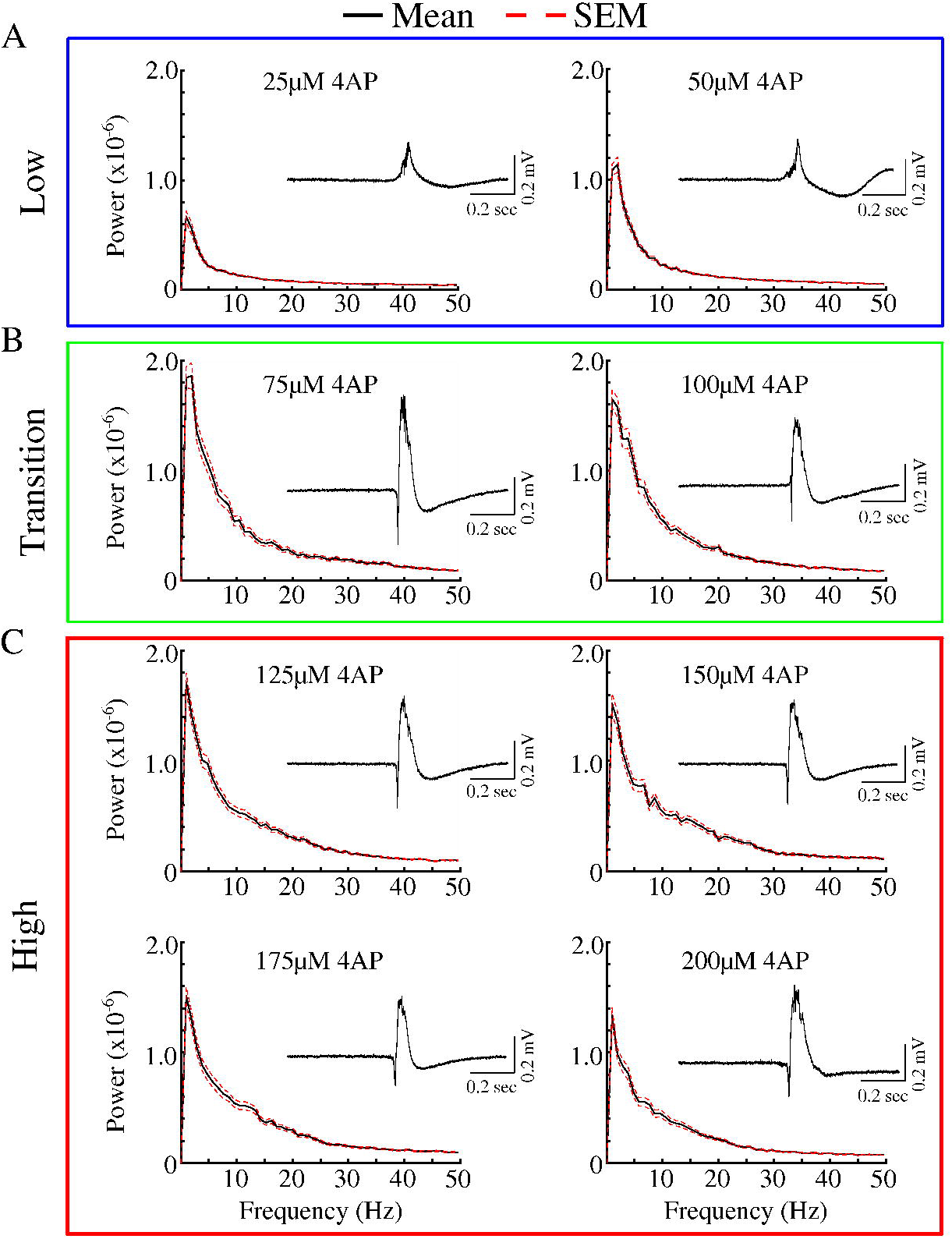
Reduction of KCC2 activity prevents concentration-dependent differences in epileptiform spectrograms. Averaged spectrograms of activity induced by different concentrations of 4AP and 10 μM VU0240551. The mean is plotted in black solid line and standard error (SEM) in red broken line. **A,** low 4AP concentrations. **B,** transition concentrations. **C,** high concentrations. Insets are of single representative epileptiform bursts elicited by the corresponding 4AP concentration. Since application of the KCC2 co-transporter antagonist resulted in overall low frequency epileptiform activity for all 4AP concentrations tested, we hypothesized that this trend may be reflected in the inter-arrival times of the epileptiform bursts. To this end, we computed the inter-event interval for different 4AP concentrations for both 4AP only and 4AP + VU0240551 conditions. As shown in figure 5A, increasing 4AP concentrations resulted in shorter inter-event intervals for both conditions (i.e. with and without VU0240551). In agreement with our hypothesis, reduction of KCC2 activity increased inter-event intervals (compare fig 5A red and black lines). For the condition of 4AP only, we observed higher event frequencies as compared to the condition of reduced KCC2 activity (fig 5B). The largest differences in event frequency was observed for lower concentrations (compare fig 5B red and black). As the concentration of 4AP increased, the frequency of events observed in the 4AP + VU0240551 condition increased to levels similar to that seen in the 4AP only condition.

**Fig 5.**
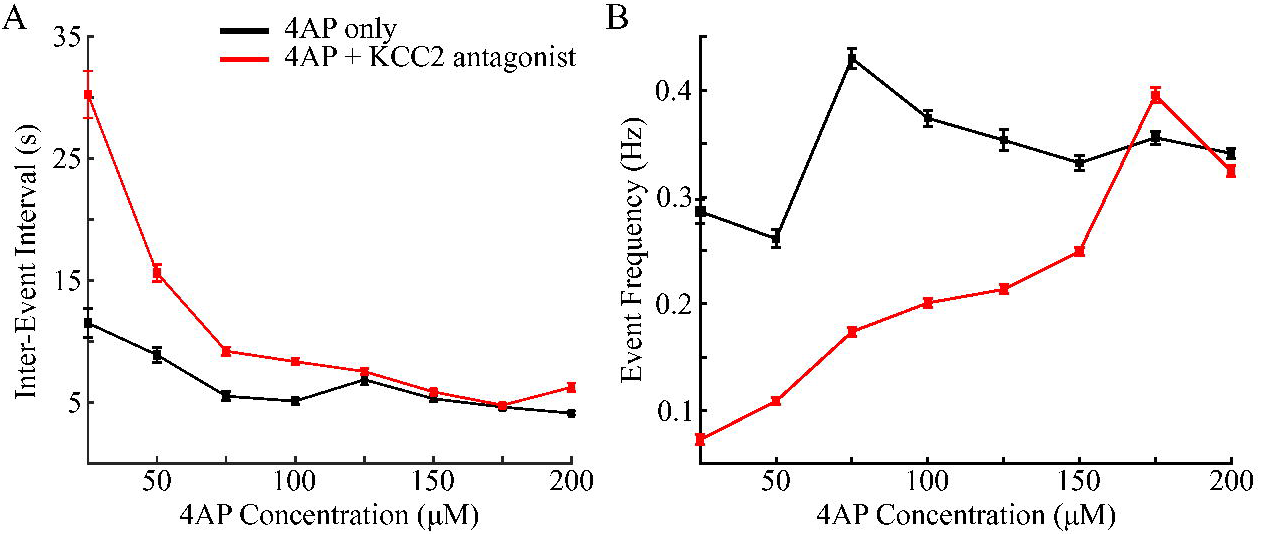
Differences in inter-event intervals and event frequency of epileptiform events. **A,** averaged inter-event intervals for difference 4AP concentrations. **B,** averaged event frequency of epileptiform events per 4AP concentration. Black indicated results for 4AP only, red for 4AP + 10 μM VU0240551.

## Discussion

K^+^-channels regulate the intrinsic excitability of neurons and maintain the repolarization phase of action potential (1-6, 30-32). Specifically, the I_A_ and I_D_ seem to account for nearly all of the K^+^-current during this repolarization phase in principal neurons (3). Due to the roles of these two currents in the regulation of neuronal excitability, they have been the focus of decades of intense study with the hope of gaining a better understanding of the mechanisms leading to the development of epilepsies (8, 13-15, 20, 26). In this new study, we tested the hypothesis that effect of I_A_ and I_D_ channel antagonist 4-aminopyridine (4AP) is strongly concentration-dependent and different 4AP concentrations are characterized by qualitatively different spatiotemporal patterns of epileptiform activity in acute mouse hippocampal brain slices. Our *in vitro* data suggests that the spatiotemporal propagation, underlying frequency components, and inter-event intervals of 4AP-induced epileptiform activity all vary with concentration. Furthermore, high concentrations of 4AP lead to appearance of the high frequency component in epileptiform events which is dependent on the activation of the KCC2 co-transporter.

We found that for all concentrations used in our study, epileptiform activity originated in the CA3 hippocampal subfield before propagating to CA1 and DG. This result is similar to previously reported findings that the CA3 is the initiator of epileptiform activity induced by elevated K^+^ (29, 33). The local architecture of the CA3 makes it a prime candidate for seizure generation. Indeed, the pyramidal neurons within the CA3 exhibit a high degree of recurrent connectivity (34). The differences in the spatio-temporal patterns of epileptiform activity arise during the application of “transition” 4AP concentrations (75 – 100 μM) and more so at high concentrations (125 – 200 μM). Low 4AP concentrations (25 – 50 μM) generate longer lasting epileptiform activity as compared to those induced by transitional and high concentrations. This may be due to altered excitability of hippocampal pyramidal neurons in response to reduced I_A_ and I_D_. Previous studies in cultured pyramidal neurons showed that the elimination of I_A_, by expression of K_v_4.2 dominant negative mutant, resulted in increased neuronal excitability to low-amplitude current stimulation (5). Interestingly, large-amplitude current injections generated transient spiking before the neuron ceased spiking activity (5). This reduction of prolonged spiking as a result of I_A_ reduction may explain the reduced durations of epileptiform activity seen in at higher 4AP concentrations.

One of the more prominent concentration-dependent features we observed was the changes in the frequency components of the induced epileptiform activity. For low concentrations of 4AP (25 – 50 μM), the epileptiform activity was dominated by a low frequency component (∼5 Hz). As the concentration was increased towards 200 μM an additional high frequency component (∼ 10 Hz) appeared. These changes in frequency were accompanied by overall reduction in the inter-event interval and increases in the number of events. This, along with the other observed concentration-dependent differences may be due to the relative differences in 4AP sensitivity exhibited by I_A_ and I_D_. I_D_ has been shown to be much more sensitive to 4AP as compared to I_A_ (3). As such, it is likely that the I_D_ is selectively blocked at the low concentrations used in this study and is responsible for the low frequency epileptiform activity. As we increase the 4AP concentration, 4AP may also block the I_A_, in addition to I_D_. This effect may explain the interesting transition observed for 75 – 100 μM 4AP. Additional experiments are needed to determine the relative contributions of these two currents in the development of epileptiform activity.

Recent work has pointed to the role of the KCC2 co-transporter in the development of 4AP-induced seizure (15, 17-19, 22). KCC2-dependent efflux and accumulation of extracellular K^+^ may drive transitions to a seizure-state by initiating a feedback loop in which elevated K^+^ leads to network depolarization, which in turn leads to further accumulation of extracellular K^+^ (17, 23, 35-40). Previous studies have shown that 4AP-induced ictal, seizure-like activity can be abolished in response to reduced KCC2 co-transporter activity (18). With this in mind, we tested whether KCC2 co-transporter inactivation could prevent the changes in epileptiform activity induced at high 4AP concentrations. Our results showed that the high frequency component of the epileptiform activity was abolished in response to KCC2 antagonist VU0240551. Additionally, the number of epileptiform events was no longer dependent on 4AP concentration, and the inter-event interval was increased. These data may point to differences in the mechanisms by which I_A_ and I_D_ reduction leads to seizure generation, with the former involving KCC2-dependent K^+^ efflux and accumulation.

In sum, our study revealed complex effect of 4AP application, possibly involving several complimentary biophysical mechanisms dependent on both ionic currents and exchangers. It further suggests that the relative contribution of these mechanisms to epileptiform activity and the overall effect of 4AP application may vary with the 4AP concentration.

## Acknowledgements

This study was supported by NIH (R01 DC012943), MURI (N000141612829), UC MRPI (MR-15-328909). Oscar C González received support from the NSF Graduate Research Fellowship under grant DGE-1326120.

